# A pilot study of a novel molecular host response assay to diagnose infection in patients after high-risk gastro-intestinal surgery

**DOI:** 10.1101/505248

**Authors:** Diana M. Verboom, Marlies E. Koster-Brouwer, Jelle P. Ruurda, Richard van Hillegersberg, Mark I. van Berge Henegouwen, Suzan S. Gisbertz, Brendon P. Scicluna, Marc J.M. Bonten, Olaf L. Cremer

**Affiliations:** Julius Center for health sciences and primary care, University Medical Center Utrecht, Utrecht, the Netherlands; Department of Intensive Care, University Medical Center Utrecht, Utrecht, the Netherlands; Center for Experimental and Molecular Medicine, Amsterdam University Medical Centers, location Academic Medical Center, University of Amsterdam, Amsterdam, the Netherlands; Department of Surgery, University Medical Center Utrecht, Utrecht, the Netherlands; Department of Surgery, Amsterdam University Medical Centers, location Academic Medical Center, Amsterdam, the Netherlands; Department of Clinical Epidemiology and Biostatistics, Amsterdam University Medical Centers, location Academic Medical Center, Amsterdam, the Netherlands; Department of Medical Microbiology, University Medical Center Utrecht, Utrecht, the Netherlands

## Abstract

**Aim:** SeptiCyte LAB measures the expression of four host-response RNAs in blood to distinguish sepsis from sterile inflammation. Sequential monitoring of this assay may have diagnostic utility in patients at high risk for postoperative infectious complications.

**Methods:** In this pilot study we studied esophagectomy patients who had developed a complication within 30 days following surgery as well as a random sample of 100 uncomplicated postoperative patients. PAXgene blood samples were collected postoperatively and whenever a complication occurred. SeptiCyte scores (ranging 0-10 with increasing likelihood of infection) were compared to post-hoc physician adjudication of infection likelihood using strict definitions.

**Results:** Among 370 esophagectomy patients, 120 (32%) subjects developed a complication requiring ICU (re)admission, 63 (53%) of whom could be analyzed. Immediate postoperative SeptiCyte LAB scores were highly variable, yet similar for patients having a complicated and uncomplicated postoperative course (median score of 2.4 (IQR 1.6-3.3) versus 2.2 (IQR 1.3-3), respectively). Scores increased as complications developed, but this rise was higher for 34 subjects having confirmed infection (median difference 4.7 (IQR 4.1-5.8)) then for 12 subjects with a non-infectious complication (2.1 (IQR 0.4-3.6); p<0.0001). When 17 cases with undetermined infectious status were excluded, addition of SeptiCyte LAB to CRP resulted in improved diagnostic discrimination (AUC 0.88 (95%CI 0.77-0.99)) compared to CRP alone (AUC 0.76 (95%CI 0.61-0.91); p=0.04).

**Conclusions:** Sequential measurement of SeptiCyte LAB may have clinical utility in surgical patients at high risk of postoperative infection, but its diagnostic performance in this setting needs to be further evaluated.

## Introduction

Major gastro-intestinal surgery, such as esophageal, gastric, and pancreatic interventions, are associated with a high risk of developing postoperative complications [1, 2]. Indeed, complication risks can be as high as 33.5%, and affected patients suffer from a significantly increased length of stay and excess mortality [2, 3]. Most of the complications are of infectious origin (principally pneumonia and anastomotic leakage), which can readily lead to the development of sepsis in postoperative patients [1–3].

Timely recognition and treatment of and treatment of sepsis may improve outcome [4]. However, sepsis diagnosis is complicated by the almost universal presence of Systemic Inflammatory Response Syndrome (SIRS) in patients who just had gastrointestinal surgery [7]. Currently, sequential measurements of C-reactive protein (CRP) and procalcitonin (PCT) are most commonly used to monitor onset of postoperative infectious complications [8]. Although these biomarkers have adequate negative predictive values (ranging from 91% to 100%), their positive predictive values remain poor [9–13]. A variety of alternative biomarkers have thus been proposed for the diagnosis of sepsis, however their diagnostic accuracy is variable across settings and their clinical utility not always evident [5, 6].

Whole-blood transcriptomics-based technologies (i.e., measurement of RNA transcripts that are generated during gene expression in leukocytes) can detect rapid change when the host is exposed to infectious stress, and may therefore possibly yield an earlier diagnostic signal of sepsis than traditional protein biomarkers [14–17]. SeptiCyte™ LAB is the first RNA-based host response signature cleared by the US Food and Drug Administration for sepsis diagnosis [18]. It measures the expression of four genes (carcinoembryonic antigen-related cell adhesion molecule 4 (CEACAM4), lysosomal-associated membrane protein 1 (LAMP1), phospholipase A2 group VII (PLA2G7), and placenta-specific 8-gene protein (PLAC8)) in peripheral blood. In several evaluations SeptiCyte LAB discriminated better between critically ill patients with (overt) sepsis and a non-infectious SIRS than PCT [17, 19, 20], although the test performed less favorably in a recent cohort of difficult-to-diagnose cases of (nosocomial) sepsis after prolonged prior hospitalization [21].

We hypothesized that sequential measurement of SeptiCyte LAB could have superior diagnostic performance over established biomarkers in postoperative patients at high risk for infectious complications, as the test characterizes the host response to infection at a relatively early —and thus possibly more specific— stage [22]. To explore this idea further, we performed a pilot study in consecutive esophagectomy patients in order to 1) determine a normal range for SeptiCyte LAB measured directly following esophageal surgery, 2) evaluate temporal changes in SeptiCyte LAB scores as complications ensue in the postoperative setting, and 3) compare the ability of SeptiCyte LAB to discriminate infectious from non-infectious complications in postoperative patients to that of a more commonly used biomarker, CRP.

## Methods

### Study design

We performed a case-control analysis that was nested in the MARS (Molecular Diagnosis and Risk Stratification of Sepsis) cohort, which enrolled subjects in two Dutch university hospitals from 2011 to 2013. From this cohort we selected consecutive patients who had undergone elective esophageal resection. Ethical approval for the study was provided by the Medical Ethics Committees of both participating hospitals, and an opt-out procedure to obtain consent from patients was in place (protocol number 10-056). Blood samples for RNA analysis were collected within 24 hours of surgery and whenever a complication occurred in the ICU during the first 30 postoperative days. This complication could be either a (suspected) infection, acute kidney injury (AKI), acute respiratory distress syndrome (ARDS), acute myocardial infarction (AMI), or readmission to the ICU for any (other) reason. For the present study, we selected cases having ≥1 postoperative complication and for whom (at least) a single paired RNA sample was available. In addition, we randomly selected 100 control subjects having an uncomplicated postoperative course after esophagectomy in order to establish (a range of) normal SeptiCyte LAB scores following major surgery.

Samples were collected in 2.5 mL PAXgene blood RNA tubes and processed in accordance with predefined acceptance criteria as set by the manufacturer of the assay (Immunexpress, Seattle, WA) [18]. Tests were performed in 96-well microtiter amplification plates on an Applied Biosystems 7500 Fast Dx Real-Time PCR Instrument (Thermo Fisher Scientific, Carlsbad, CA), yielding a threshold cycle number (Ct-value) per individual gene. A score was then calculated as *(Ct_PLA2G7_ + Ct_CEACAM4_) – (Ct_PIAC8_ + Ct_LAMP1_)*. This ‘SeptiScore’ ranges from 0 to 10, and may be categorized into 4 probability bands according to the manufacturer’s specification [18]. SeptiScores ≤3.0 (band 1) indicate that sepsis is unlikely, whereas scores 3.1—4.4 (band 2), 4.5—5.9 (band 3), and >6 (band 4) represent increasing sepsis likelihoods.

### Reference test for infection

Suspected infectious events were recorded prospectively upon each occasion that antimicrobial therapy was initiated by the clinician. All patients treated for a suspected infectious event in the ICU also met SIRS criteria and had a Sequential Organ Failure Assessment (SOFA) score ≥2, thus fulfilling current Sepsis-3 definitions [24, 25]. The likelihood of infection was subsequently classified as none, possible, probable, or definite based on daily discussions with the attending team as well as a post-hoc review of all available clinical, microbiological, and radiological data collected during ICU stay by trained physicians according to predefined definitions [23]. This reference diagnosis was established without knowledge of SeptiCyte LAB results, yet observers were not blinded to CRP. However, CRP in and of itself could not lead to a diagnosis of infection in the absence of other clinical and inflammatory symptoms.

For use as a reference test in the current study, all observed complications were reclassified as infection ruled-out (patients with a post-hoc likelihood rated none, or patients who were never suspected of an infection), infection undetermined (patients with possible infection), or infection confirmed (patients with a post-hoc likelihood rated probable or definite). In case of multiple concurrent events, SeptiCyte LAB test results were related to the complication that occurred nearest in time to the moment the sample was taken.

### Statistical analysis

Immediate postoperative SeptiScores were analyzed to determine a normal range following major surgery, both in patients who would later develop a complication as well as in the 100 control subjects. Subsequently, in esophagectomy patients having a complicated postoperative course only, we performed within-patient pairwise comparisons of scores measured in samples obtained directly after surgery and at complication onset. To assess the diagnostic potential of sequential SeptiCyte LAB measurement, we focused this analysis on differences between subjects having non-infectious, undetermined and confirmed infectious complications. In addition, we compared SeptiScores to CRP concentrations measured in plasma obtained at the same time point. To this end, we standardized differences between the post-operative moment and the moment of complication onset by calculating Z-scores. We compared discriminative ability of SeptiCyte LAB and CRP using receiver operating characteristic (ROC) curves. For this latter analysis we excluded subjects having an undetermined infectious state according to physician adjudication.

Differences in categorical and continuous variables between groups were assessed using Chi-square, Wilcoxon signed rank or Mann-Whitney U tests, as appropriate. All analyses were performed in SAS Enterprise Guide 7.1 (SAS Institute, Cary, NC) and R Studio (R Studio Team (2015), Boston, MA).

## Results

During the study period, a total of 370 patients were admitted to the ICU after elective esophagectomy, of whom 120 (34%) developed a complication resulting in prolonged ICU stay or readmission to the ICU within 30 days. Among these, 74 (62%) subjects had immediate postoperative PAXgene blood samples available for analysis, and 63 (53%) also had a sample taken at complication onset (figure 1). Patients without a postoperative sample (n=46) had a shorter ICU stay (9, IQR 2-16 versus 12, IQR 7-24, p=0.04), less ICU readmissions (54% versus 97%, p<0.001), and were treated for infection less frequently (70% versus 92%, p<0.05) than patients with an available sample. However, in-hospital mortality was similar between the groups (13% versus 16%, p=0.79).

**Fig. 1.**
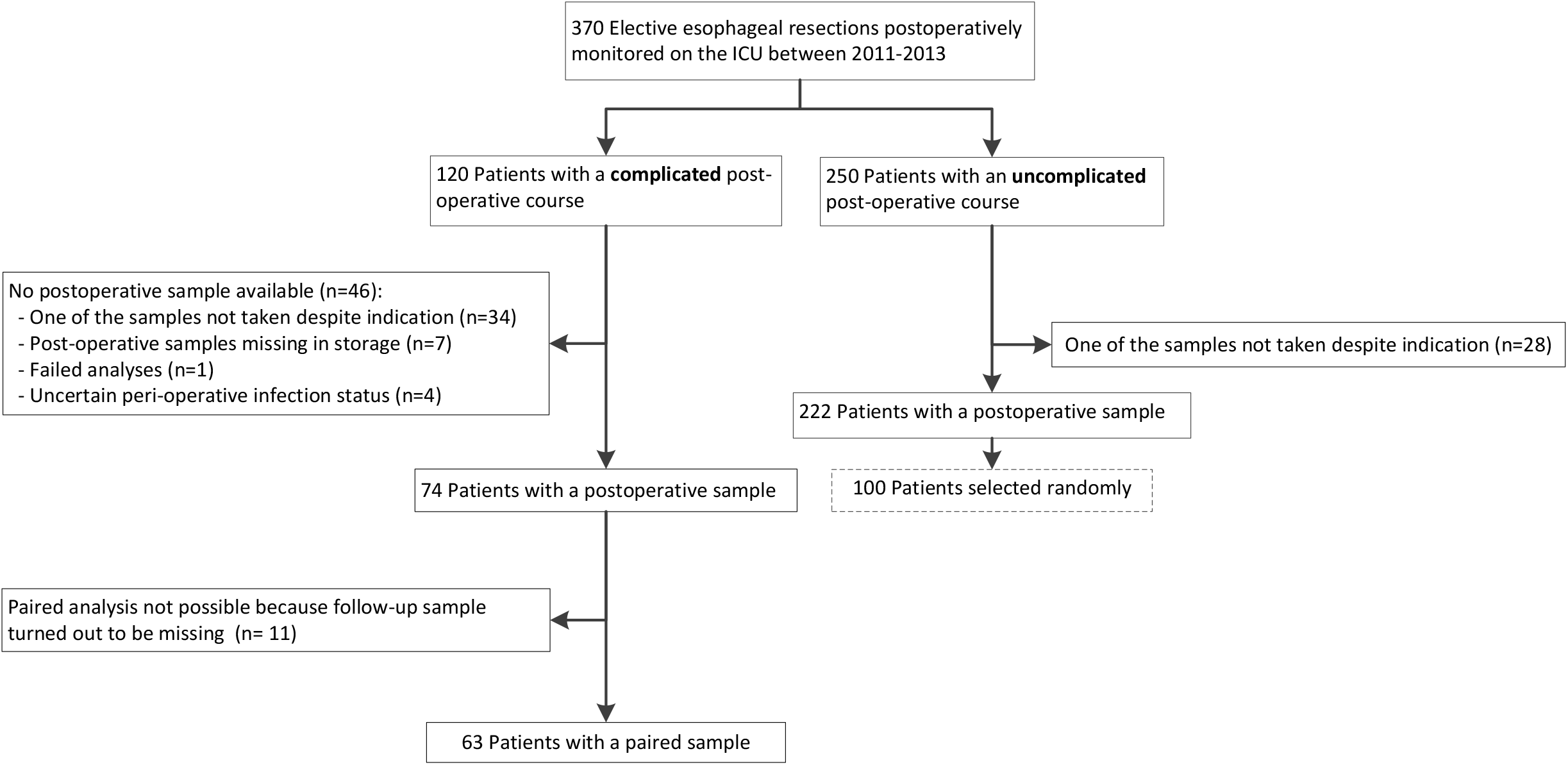
Flowchart of patient inclusion. *ICU*; Intensive Care Unit.

Immediate postoperative PAXgene samples were additionally analyzed in a random subsample (n=100) of the 250 remaining esophagectomy patients who had an uncomplicated postoperative course. These patients less frequently developed SIRS, required less vasopressors, and exhibited a trend towards lower APACHE scores and CRP levels compared to patients in the complicated cohort (table 1).

**Table 1.**
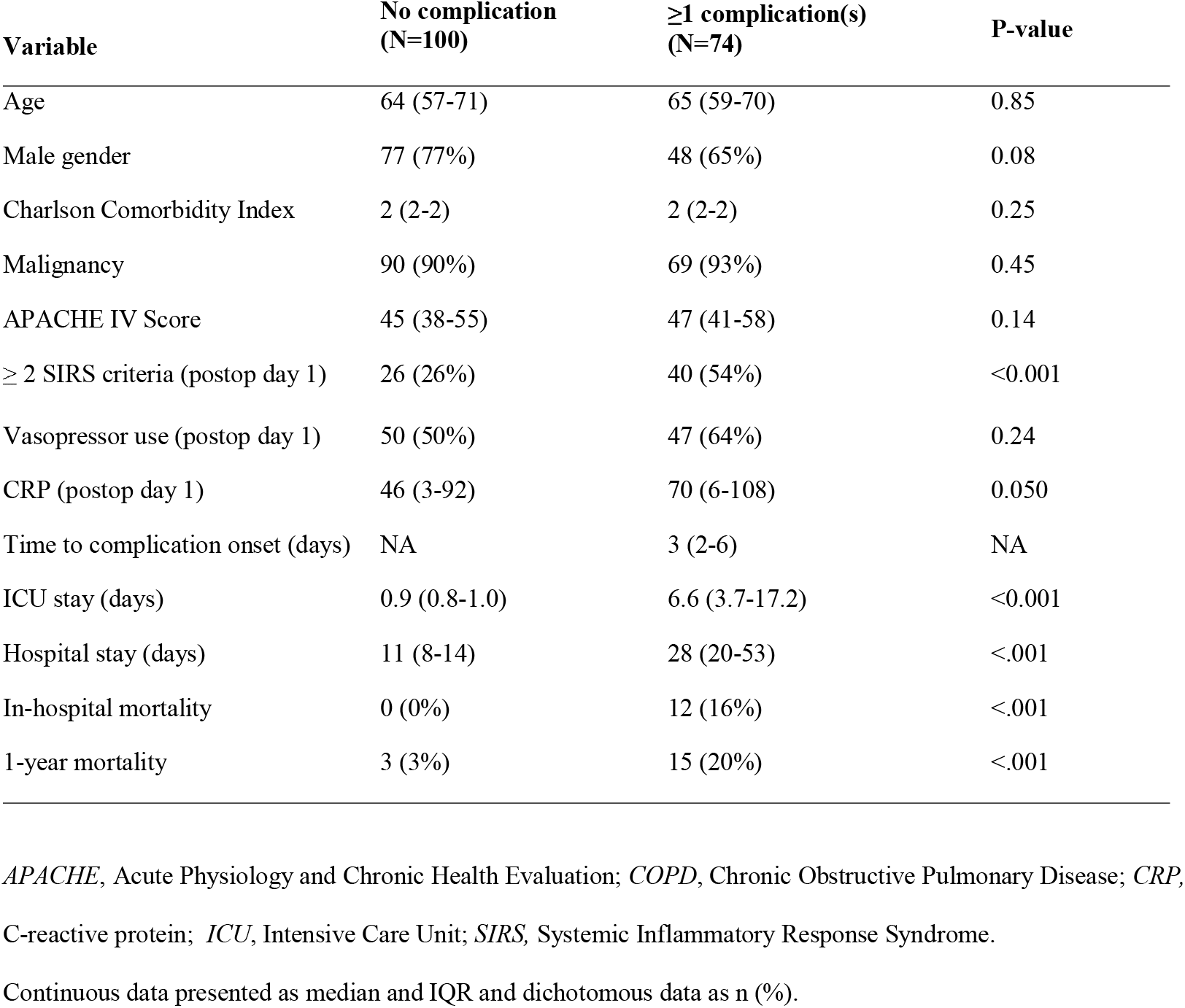
Characteristics of 174 esophagectomy patients stratified by their postoperative clinical course

Among the 63 patients with postoperative complications who were included in the pairwise analyses of SeptiCyte LAB, 34 (54%) subjects had a confirmed infection, 17 (27%) an undetermined infectious state, and 12 (19%) a non-infectious complication (i.e., 5 were empirically treated with antibiotics but classified as having no infection in retrospect, whereas 7 were never suspected of infection). Frequently observed infections included intrathoracic sources (34%; most commonly mediastinitis or pleural empyema due to anastomotic leakage) and pneumonia (24%) (table 2). Of note, multiple complications could coexist.

**Table 2.** Clinical events among 63 esophagectomy patients with ≥1 postoperative complication

| Complications | Non-infectious<br>(n=12) | Undetermined<br>(n=17) | Confirmed infection<br>(n=34) |
| --- | --- | --- | --- |
| Total number of events | 18 | 39 | 97 |
| *(Presumed) infectious source(s)* |  |  |  |
| - intrathoracic<sup>a</sup> | 0 (0%) | 6 (15%) | 24 (25%) |
| - pulmonary | 4 (22%)<sup>b</sup> | 10 (26%) | 11 (11%) |
| - abdominal | 1 (6%)<sup>b</sup> | 2 (5%) | 7 (7%) |
| - other | 0 (0%) | 0 (0%) | 2 (2%) |
| *Other (non-infectious) complication(s)* |  |  |  |
| - readmission to ICU | 10 (56%) | 16 (41%) | 33 (34%) |
| - acute respiratory distress syndrome | 1 (6%) | 3 (8%) | 13 (13%) |
| - acute kidney injury | 2 (11%) | 1 (3%) | 7 (7%) |
| - acute myocardial infarction | 0 (0%) | 1 (3%) | 0 (0%) |

ICU, Intensive Care Unit. Multiple complications could occur at the same time.
<sup>a</sup>Intrathoracic infections include mainly cases of lung empyema and mediastinitis
<sup>b</sup>Initially suspected focus of infection among patients in whom infection likelihood was later classified as 'none'

### SeptiScore distribution in the immediate postoperative setting

Among the total number of 174 analyzed patients, immediate postoperative SeptiScores were highly variable (range 0-10), with a median of 2.3 (IQR 1.4-3.1). Overall, 45 (26%) samples corresponded to probability band 2 or higher, which —in case sepsis were to be clinically suspected— would have incorrectly resulted in a “sepsis-likely” label according to the manufacturer’s specification (table 3). Median SeptiScores of patients having an uncomplicated postoperative course tended to be slightly lower than those of patients who would later develop a complication (2.2 (IQR 1.3-3) versus 2.4 (IQR 1.6-3. 3)), although this difference did not reach statistical significance (p= 0.14).

**Table 3.** Number of samples by SeptiCyte LAB probability band and reference test

| SeptiCyte LAB probability band* | Interpretation | Reference test |  |  |  |
| --- | --- | --- | --- | --- | --- |
|  |  | Postoperative<br>(n=174) | Non-infectious<br>complication<br>(n= 12) | Undetermined<br>(n=17) | Confirmed<br>infection (n=34) |
| Band 1 (SeptiScore 0-3) | Sepsis unlikely | 129 (74) | 1 (8) | 1 (6) | 0 (0) |
| Band 2 (SeptiScore 3.1-4.4) | Sepsis likely | 28 (16) | 2 (17) | 1 (6) | 0 (0) |
| Band 3 (SeptiScore 4.5-5.9) | Sepsis likely | 14 (8) | 6 (50) | 4 (24) | 7 (21) |
| Band 4 (SeptiScore 6-10) | Sepsis likely | 3 (17) | 3 (25) | 11 (65) | 27 (79) |

Presented as n (%). Percentages are column percentages. Reference test presented by post-hoc likelihood.
\*Higher SeptiCyte LAB probability bands indicate increased likelihood of sepsis

### Temporal changes in SeptiScore

Figure 2 shows SeptiScores as measured immediately after surgery and at the time of complication onset for the 63 patients in whom paired samples were available. Median lag time between surgery and the development of a complication was 5 (IQR 3-9) days. However, it should be noted that 8 (14%) of the repeat samples were already taken within 2 days of surgery. SeptiScores increased in all patients who developed a complicated disease course after surgery, but this rise was more pronounced in those developing infection versus another complication (median score differences 2.1 (IQR 0.4-3.6), 4 (IQR 2.5-5), and 4.7 (IQR 4.1-5.8) for subjects having a non-infectious, undetermined, and confirmed infectious event, respectively; p<0.0001 (table 4).

**Fig. 2.**
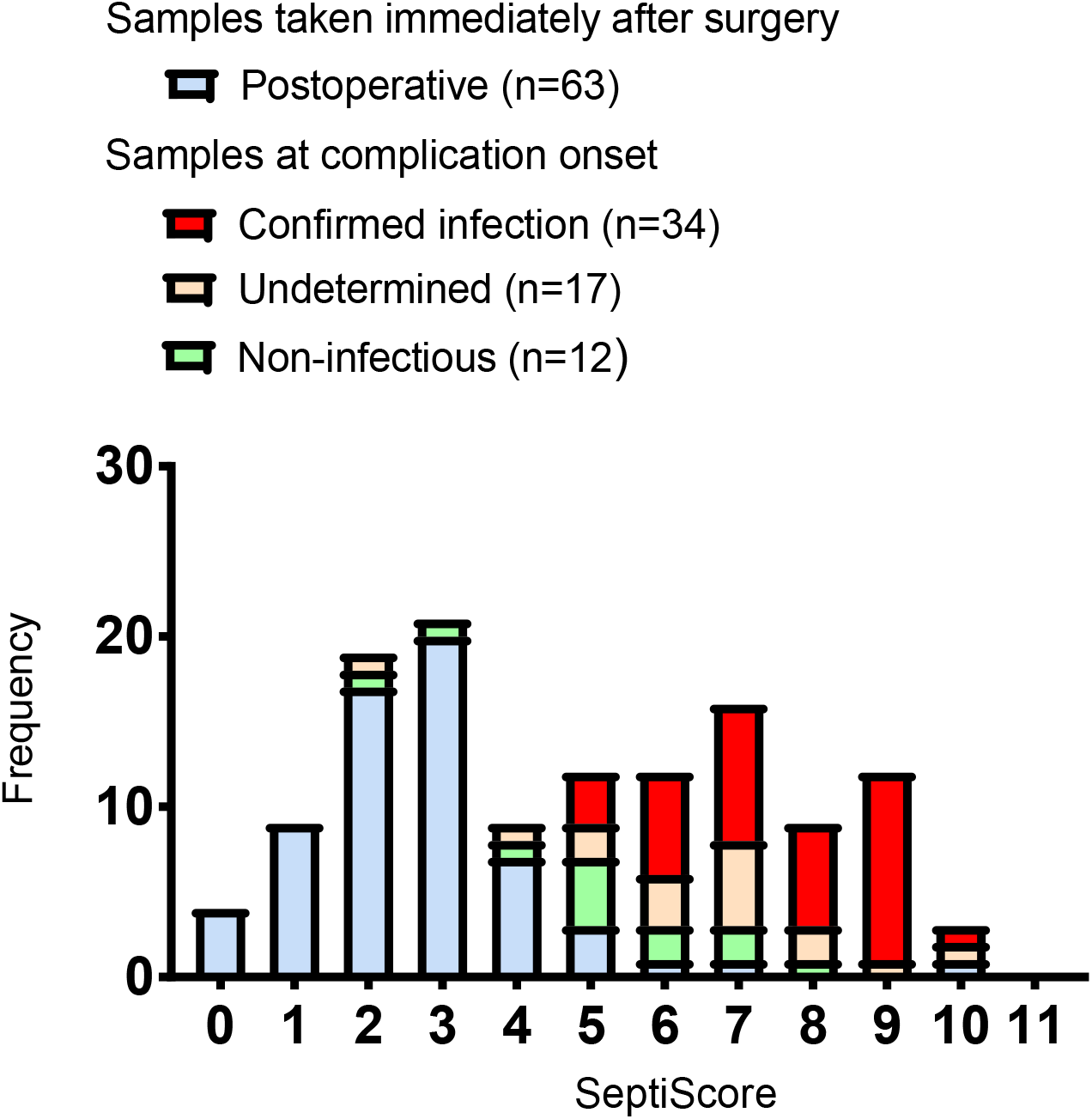
SeptiScores among 63 esophagectomy patients with ≥1 postoperative complication. Frequencies represent the number of patients.

**Table 4.** SeptiScore and CRP in infectious complications and non-infectious complications in 63 patients

|  | **Non-infectious**<br>**(n=12)**<br>**Median (IQR)** | **Undetermined**<br>**(n=17)**<br>**Median (IQR)** | **Confirmed infection**<br>**(n=34)**<br>**Median (IQR)** | **p-value<sup>a</sup>** |
| --- | --- | --- | --- | --- |
| SeptiScore |  |  |  |  |
| - immediate postop sample | 2.7 (1.4-4.6) | 2.7 (2.2-2.9) | 2.5 (1.6-3.4) | - |
| - complication onset sample | 5.3 (4.2-6.1) | 6.8 (5.5-7.4) | 7.6 (6.3-8.7) | 0.0002 |
| - delta score | 2.1 (0.4-3.6)* | 4.0 (2.5-5)* | 4.7 (4.1-5.8)* | <0.0001 |
| CRP concentration (mmol/L) |  |  |  |  |
| - immediate postop sample | 87 (46-132) | 70 (4-108) | 60 (9-102) | - |
| - complication onset sample | 184 (128-275) | 257 (181-320) | 315(194-386) | 0.0083 |
| - delta concentration | 124 (53-197)* | 185 (107-279)* | 206 (162-312)* | 0.0043 |

*CRP*, C-reactive protein. Delta is the difference in SeptiScore and CRP between the postoperative moment and the moment when a complication occurred.
<sup>a</sup>Mann whitney U Test comparing non-infectious complication to confirmed infection.
\*Wilcoxon signed rank test for the delta $p < 0.05$ , meaning that the delta was significantly different from 0.

### Discriminative ability of SeptiCyte LAB

Standardized differences (expressed as Z-scores) between samples collected immediately after surgery and at complication onset revealed a greater increase in patients with confirmed infections than in other patients for both SeptiCyte LAB and CRP (figure 3). However, among the 34 patients with confirmed infection, the observed standardized increase in SeptiScore was more pronounced than that in CRP, although this difference did not reach statistical significance (median Z-score 0.28 versus 0.08, p=0.08).

**Fig. 3.**
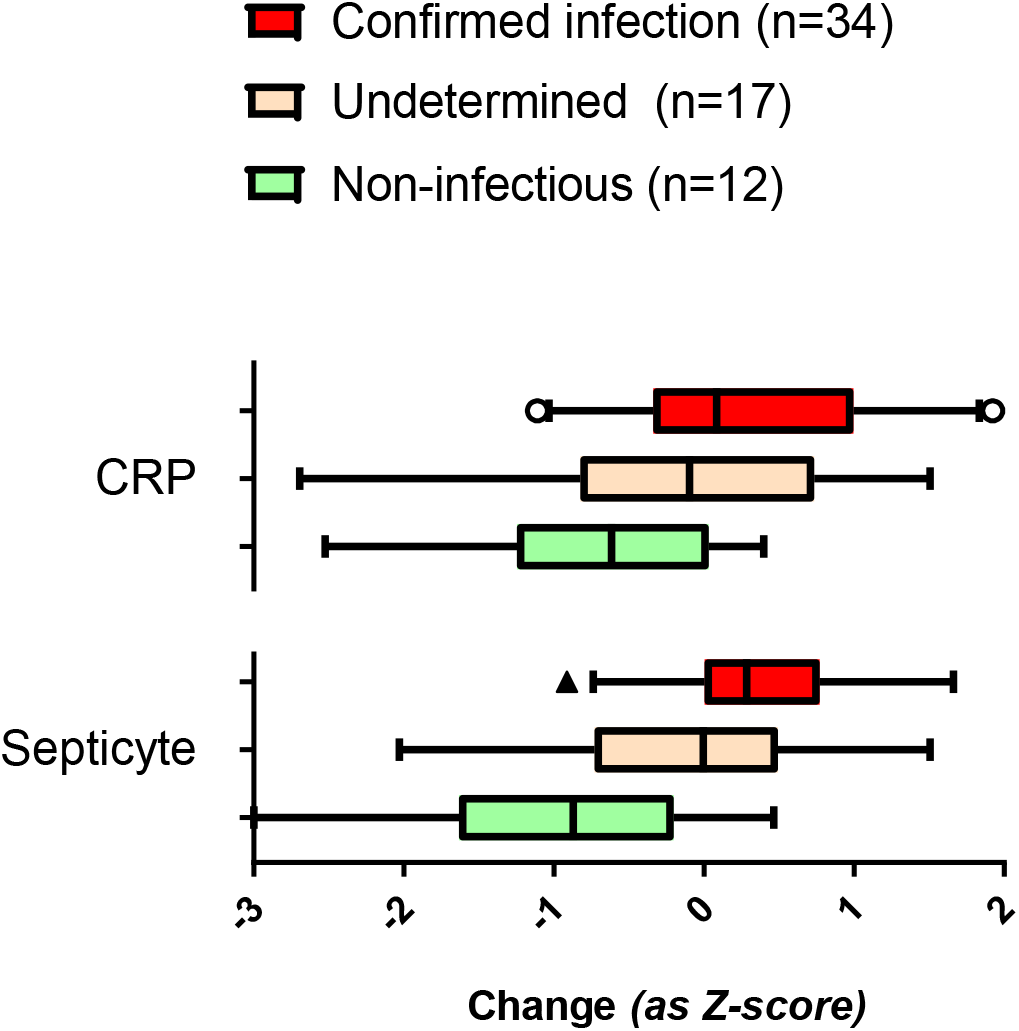
Temporal changes in SeptiScores and CRP levels among 63 esophagectomy patients developing a postoperative complication. *CRP*, C-reactive protein. CRP concentration and SeptiScores were measured in the immediate postoperative sample and the sample taken at complication onset. Subsequently, observed differences between both time points were transformed into a standardized z-score (z=x-μ/σ), having a mean value of 0 and standard deviation of 1. Boxes show median standardized differences with IQR, whiskers show 5-95^th^ percentiles.

In a direct comparison of patients developing a confirmed infectious (n=34) and definite non-infectious complication (n=12), ROC analysis yielded an AUC of 0.87 (95%CI 0.760.98) for SeptiCyte LAB, compared to an AUC of 0.76 (95%CI 0.61-0.91) for CRP (p=0.14). Adding SeptiCyte LAB to CRP resulted in improved diagnostic discrimination (AUC 0.88 (95%CI 0.77-0.99), p=0.04).

## Discussion

This pilot study explored whether temporal changes in SeptiCyte LAB could be used to help diagnose infectious complications after esophageal surgery. Although SeptiScores varied widely between individuals, median scores immediately after surgery were comparable between subjects who went on to have either an eventful or uncomplicated subsequent postoperative course. However, the increase of SeptiScore over time was greater in patients developing postoperative infections than in those with other complications. Furthermore, this appeared to be more pronounced than the simultaneous rise in CRP observed in these patients.

SeptiCyte LAB was originally developed to help diagnose infection in critically ill patients presenting to an ICU with SIRS. Previous studies evaluating its diagnostic performance in this setting have reported variable discriminative ability for SeptiCyte LAB, with AUC’s ranging from 0.73 (0.68-0.79) to 0.99 (95%CI 0.96-1.00) in different cohorts [17, 19–21, 26, 27]. In particular, specificity was lower in patients presenting with suspected pneumonia as well as in those who had already been subjected to a prolonged clinical course in hospital prior to ICU admission [21, 26]. In the current study we observed favorable discrimination (AUC of 0.87 (95%CI 0.76-0.98), albeit after exclusion of patients having an uncertain infectious state.

Observed variations in diagnostic performance are most likely explained by differences in study size, clinical setting, and distribution of underlying infectious etiologies. The optimal intended use scenario for SeptiCyte LAB therefore requires further exploration. Our data suggest that adding SeptiCyte LAB to CRP may improve diagnostic discrimination in patients following major surgery [9–12]. However, any possible use of SeptiCyte LAB for routine screening of such postoperative patients will require careful evaluation before it can be recommended, as the test will probably be more expensive than alternative biomarkers that are routinely available. As CRP has good negative predictive value, a stepped approach therefore seems reasonable (i.e., by measuring SeptiCyte LAB only in patients with increased CRP). Clinical utility and cost-effectiveness of such strategy must be evaluated in a randomized controlled diagnostic trial.

Our pilot study has several limitations related to its relatively small sample size, which limits statistical power, as well as the unavailability of paired PAXgene samples in almost half the target population. In addition, not all study patients presented to the ICU having a true diagnostic dilemma regarding the presence or absence of infection, which precludes final conclusions regarding diagnostic accuracy. Although our pilot series aimed to explore the potential utility of sequential measurement of SeptiCyte LAB in the postoperative setting, we did not actually assess this biomarker on a daily basis. Thus, our diagnostic findings cannot be directly extrapolated to a setting of sequential daily monitoring. Also, for the prompt initiation of PAXgene specimen collection researchers were dependent on the clinical recognition of complications by attending physicians, which may have led to between-patient variability in timing of samples. Finally, even though the post-hoc likelihood of all suspected infections was carefully adjudicated by trained physician-observers according to standardized definitions, diagnostic misclassification cannot be ruled out [23]. However, patients with the greatest uncertainty regarding their reference diagnosis (i.e., those with an undetermined infectious status), were excluded from most comparative analyses, as has been done before in similar studies[17, 21, 28].

In conclusion, SeptiCyte LAB may have clinical utility in surgical patients at high risk for postoperative infection. However, its diagnostic value in this setting has to be further evaluated in prospective studies using sequential sampling both in patients merely at risk for developing postoperative infections, as well as in those with clinically suspected sepsis onset after major surgery.

## Compliance with ethical standards

Ethical approval for the study was provided by the Medical Ethics Committees of both participating hospitals, and an opt-out procedure to obtain consent from eligible patients was in place (protocol number 10-056). This study was therefore performed in accordance with the ethical standards laid down in the 1964 Declaration of Helsinki and its later amendments. The study was registered at clinicaltrials.gov (identifier NCT01905033).

## Acknowledgments

We thank the MARS consortium, including the participating ICUs and research physicians of the two medical centers for their help in data acquisition. We especially thank Tom van der Poll and Kirsten van de Groep, for their contributions to this manuscript.

